# Evaluating Protein Language Model Embeddings for Structural Similarity in the Protein-Sequence Twilight Zone

**DOI:** 10.64898/2026.07.31.742099

**Authors:** Shivaram Danwada, Priscilla Udomprasert, Nilanjana Raychawdhary, Cheryl D Seals, Lei Wu, Sutanu Bhattacharya

## Abstract

Evaluating protein sequence similarity remains challenging in the protein-sequence twilight zone (20–35% sequence identity), where traditional methods often fail. In this study, we evaluate whether mean-pooled embeddings from four protein language models: ESM-1b, ESM-2, ProtT5, and ProstT5 can estimate pairwise structural similarity without performing sequence alignment. The benchmark dataset includes 20,445 PISCES protein pairs with sequence identity ≤30%, representing the protein-sequence twilight zone, with TM-align-derived TM_min_ used as the structural ground truth. Protein embeddings are compared using cosine similarity, Euclidean- and Manhattan-derived similarities, an RBF kernel, and dot product. Among these similarity metrics, cosine similarity performs best across all four models. Moreover, ProstT5 achieves the highest Spearman correlation with TM_min_, followed by ESM-2, ProtT5, and ESM-1b, while all four PLMs outperform BLASTP overall. Furthermore, the advantage of PLM embeddings is most pronounced for protein pairs with the lowest sequence identity. ProstT5 also provides the best discrimination between structurally similar and dissimilar protein pairs. Moreover, it offers a favorable balance between similarity performance and the computational requirements of residue-level embedding generation and storage. Overall, these findings support PLM embeddings as an effective alignment-free approach for detecting structural relationships among proteins in the twilight zone.

## I. INTRODUCTION

**T**HE rapid growth of protein-sequence databases [1], [2] allows recent advances in natural language processing (NLP) to be transferred to protein representation learning [3], [4]. Protein language models (PLMs) [5]–[8] treat amino acids or residues as tokens and learn statistical regularities from large collections of naturally occurring sequences. Similar to transformer-based language models developed for NLP [9], [10], many PLMs use a self-supervised learning objective in which individual residues or contiguous residue spans from protein sequences extracted from large protein-sequence databases are masked and then reconstructed from the remaining sequence context [11]. Successful reconstruction requires the model to determine which positions in the input provide relevant contextual information. The self-attention mechanism of the transformer architecture supports this process by assigning context-dependent weights to residues throughout the sequence. During pretraining, the model consequently builds high-dimensional representations of individual residues and entire protein sequences [11]. Because these representations are learned without labels for a specific biological task, they can subsequently be used directly in zero-shot settings or adapted to a variety of down-stream applications.

Representations derived from PLMs are useful for a wide range of protein-prediction problems, including secondary-structure prediction [7], subcellular-localization prediction [7], [12], inference of evolutionary relationships [13], and classification at the family and superfamily levels [14], [15]. More importantly, several studies report that PLMs can learn structural information while trained only on amino-acid sequences [5], [7], [16], [17]. For example, TAPE [16] demonstrates that pretrained sequence representations support secondary-structure prediction, remote-homology detection, and protein-engineering tasks. ESM-1b [17] representations contain signals associated with secondary structure, long-range residue contacts, mutational effects, and homology. The ESM-2 family [18] extends this line of work from millions to billions of parameters and provides the sequence representations used by ESMFold for atomic-level structure prediction. Although sequence-only language models do not replace structure-prediction systems such as AlphaFold2 [19], their performance indicates that large-scale sequence pretraining recovers substantial information about the structural constraints acting on proteins.

The view that PLMs acquire structure-related information is biologically plausible because their training corpora contain the outcomes of evolutionary selection across a vast region of protein-sequence space. Protein function imposes constraints on three-dimensional conformation, and the requirements of a stable and functional conformation restrict the amino-acid substitutions that are tolerated during evolution [20]. The resulting selective pressures are therefore transmitted from function through structure to sequence. Function typically changes more slowly than structure, while protein structure remains substantially more conserved than primary sequence [21]. A model trained on millions of evolutionarily sampled sequences may consequently recover statistical patterns that reflect both local biochemical preferences and global structural constraints.

Structural and functional relationships are usually recognizable above approximately 30% sequence identity ≤( 30%) [22]–[29], but below this threshold direct sequence comparison becomes increasingly unreliable even when proteins retain related folds or functions [29]–[32]. This low-identity regime, known as the protein-sequence twilight zone, therefore provides a stringent setting for testing whether PLMs encode structure-relevant information beyond obvious sequence similarity.

However, it remains unclear whether sequence-level PLM embeddings can serve directly as reliable measures of pairwise structural similarity in the twilight zone, and which combination of PLM and similarity metric most effectively preserves structural relationships. Existing studies largely evaluate PLMs through remote-homology classification [4], [33], residue-level embedding alignment [3], [34]–[37], or end-to-end structure prediction [18] rather than through continuous pairwise structural-similarity scoring across multiple models and embedding-space metrics.

To address this gap, we examine this question in a zero-shot setting, using each pretrained PLM only as a fixed feature extractor and asking whether similarity between mean-pooled protein embeddings tracks experimentally determined structural similarity. The benchmark is restricted to protein pairs with no more than 30% sequence identity and is further divided into 0%≤ *x*≤ 10%, 10% < *x*≤ 20%, and 20% < *x* ≤30% intervals to determine whether performance remains stable across the twilight zone.

We evaluate four representative transformer-based PLMs — ESM-1b [17], ESM-2 [18], ProtT5 [7], and ProstT5 [38] — covering encoder-only and T5-based architectures as well as sequence-only and structure-aware pretraining. For each protein pair, the sequence-level embeddings are compared using cosine similarity, Euclidean-derived similarity, an RBF kernel, Manhattan-derived similarity, and dot product, since different metrics capture different aspects of embedding geometry and may behave differently across models. BLASTP [39], [40], a traditional protein sequence alignment tool, serves as the baseline that does not use embeddings. Overall, the results show that PLM embeddings capture meaningful structural similarity throughout the twilight zone and consistently outperform BLASTP. Among the evaluated models, ProstT5 provides the strongest structure-aware representations, while cosine similarity emerges as the most reliable metric across architectures. All our code is made publicly available at https://github.com/sutanubh1/RemoteProtBench.

The remainder of this paper is organised as follows. Section II describes the methods, Section III presents and discusses the results, and Section IV concludes the paper and outlines directions for future work.

## II. METHODS

### A. OVERVIEW OF THE PIPELINE

As shown in **Fig. 1**, given an input pair of protein sequences, *P*_1_ and *P*_2_, with lengths *L*_1_ and *L*_2_, respectively, each sequence is processed independently by a pretrained PLM. The PLM generates residue-level embeddings:

**FIGURE 1.**
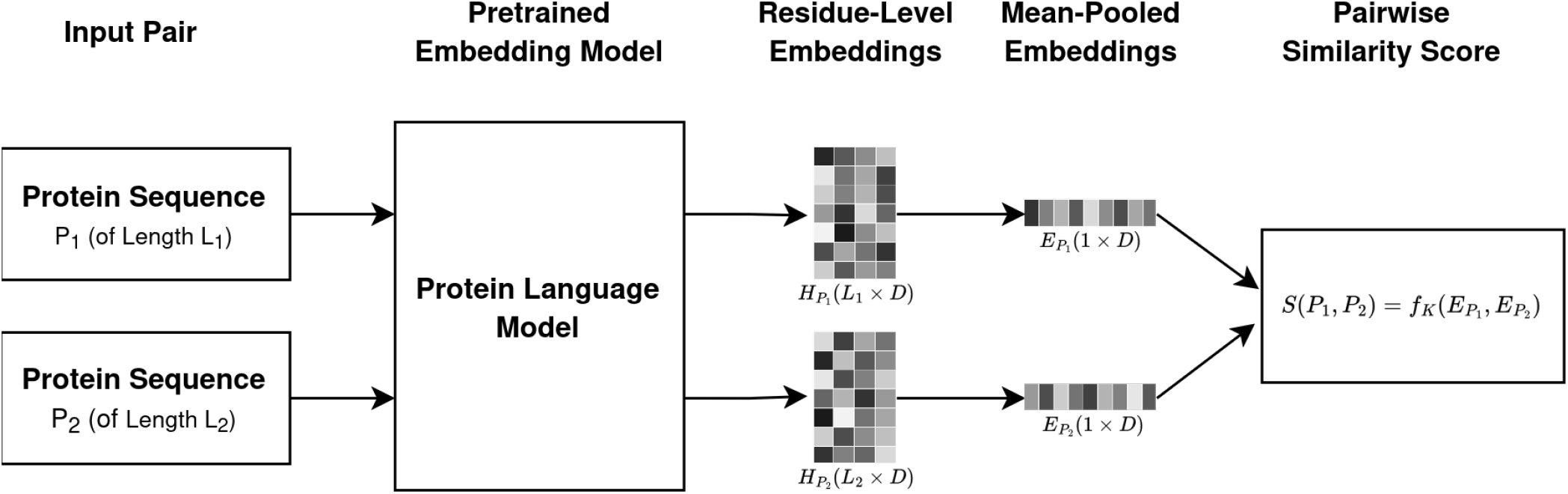
Overview of the embedding-based pipeline. Two input protein sequences, *P*_1_ and *P*_2_, of lengths *L*_1_ and *L*_2_, are independently processed by the same pretrained PLM to generate residue-level embeddings, H_*P*_ of dimension (*L*_1_× *D*) and H_*P*_ of dimension (*L*_2_ × *D*). D denotes the embedding dimension of the PLM. Mean pooling across residue positions produces fixed-length protein representations, 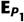 and 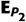, each of dimension 1 *D*. A pairwise similarity metric *f*_*k*_ is then applied to the pooled embeddings to obtain the alignment-free similarity score, *S*(*P*_1_, *P*_2_).

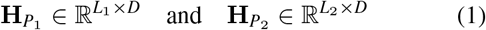

where *D* denotes the embedding dimension of the model, and 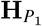 of dimension (*L*_1_ × *D*) and 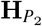 of dimension (*L*_2_ × *D*) are residue-level embeddings for *P*_1_ and *P*_2_, respectively. After removing model-specific special and padding tokens, the sequence-level representations of *P*_1_ and *P*_2_ are obtained by mean pooling the residue-level embeddings across the sequence length as follows:

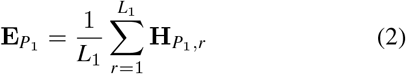

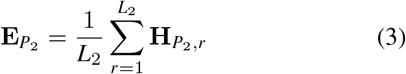

where 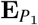 and 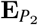 are the sequence-level representations of *P*_1_ and *P*_2_, respectively. Mean pooling produces one *D*- dimensional vector for each protein, irrespective of the original sequence length. Finally, the pairwise similarity score between the two proteins is calculated by applying a selected similarity metric *f*_*k*_ to the mean-pooled embeddings as follows:

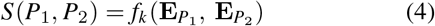

### B. SELECTED PROTEIN LANGUAGE MODELS

We evaluate four representative transformer-based PLMs: ESM-1b [17], ESM-2 [18], ProtT5 [7], and ProstT5 [38]. The models span encoder-only and encoder–decoder architectures, differ in parameter scale and embedding dimensionality, and include both sequence-only and structure-aware pretraining strategies. **Table 1** summarizes the key properties of each model, which are described below.

**TABLE 1.**
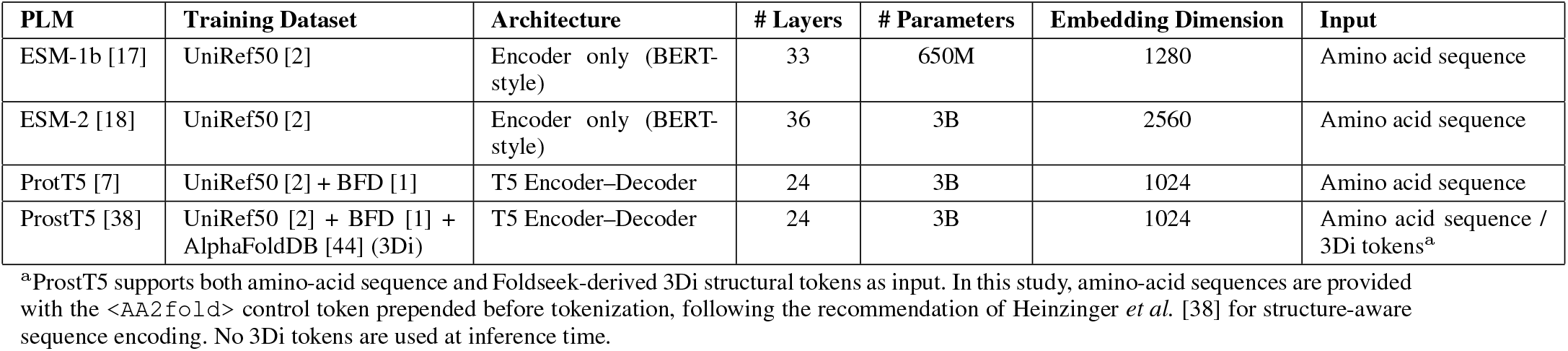
Protein language models (PLMs) evaluated in this study.

| PLM | Training Dataset | Architecture | # Layers | # Parameters | Embedding Dimension | Input |
| --- | --- | --- | --- | --- | --- | --- |
| ESM-1b [17] | UniRef50 [2] | Encoder only (BERT-style) | 33 | 650M | 1280 | Amino acid sequence |
| ESM-2 [18] | UniRef50 [2] | Encoder only (BERT-style) | 36 | 3B | 2560 | Amino acid sequence |
| ProtT5 [7] | UniRef50 [2] + BFD [1] | T5 Encoder–Decoder | 24 | 3B | 1024 | Amino acid sequence |
| ProstT5 [38] | UniRef50 [2] + BFD [1] + AlphaFoldDB [44] (3Di) | T5 Encoder–Decoder | 24 | 3B | 1024 | Amino acid sequence / 3Di tokens<sup>a</sup> |

<sup>a</sup>ProstT5 supports both amino-acid sequence and Foldseek-derived 3Di structural tokens as input. In this study, amino-acid sequences are provided with the <AA2fold> control token prepended before tokenization, following the recommendation of Heininger *et al.* [38] for structure-aware sequence encoding. No 3Di tokens are used at inference time.

#### 1) ESM-1b

ESM-1b [17] is a 33-layer bidirectional transformer encoder with approximately 650 million parameters and a residue-level embedding dimension of 1,280. It is pretrained on evolutionary-scale protein-sequence data using masked language modeling, in which selected amino-acid tokens are reconstructed from their surrounding sequence context. For a protein of length *L*, ESM-1b produces a residue-level representation **H**_*P*_∈ ℝ^*L*×1280^after beginning-of-sequence and end-of-sequence tokens are removed. Because the model has a finite input-context length, sequences longer than 1,022 residues are divided into overlapping windows, embedded separately, and recombined to recover a complete residue-level representation. ESM-1b is included because it is among the first large PLMs to demonstrate that structural, evolutionary, and functional information emerges from sequence-only self-supervised pretraining.

#### 2) ESM-2

ESM-2 [18] extends the ESM framework through architectural refinements, larger training collections, and model sizes ranging from millions to billions of parameters. We use the 3-billion-parameter variant (esm2_t36_3B_UR50D), which contains 36 transformer layers and produces a 2,560-dimensional representation for each residue. For a sequence of length *L*, the extracted embedding matrix has dimensions **H**_*P*_ ∈ ℝ^*L*×2560^after beginning-of-sequence and end-of-sequence tokens are excluded. Representations are obtained from transformer layer 33, following the default ESM-family extraction configuration used by the embedding-based alignment toolkit [34]. As with ESM-1b, sequences exceeding 1,022 residues are processed through overlapping windows and reconstructed into complete residue-level embeddings. ESM-2 is trained using masked-residue prediction on large UniRef-derived sequence collections and provides the representations used by the ESMFold structure-prediction framework.

#### 3) ProtT5

ProtT5 [7] belongs to the T5 encoder–decoder family and is developed as part of the ProtTrans framework [7]. It is pretrained on large protein-sequence collections derived from UniRef50 [2] and the Big Fantastic Database (BFD) [1], which contains extensive genomic and metagenomic sequence diversity. The complete model contains approximately 3 billion parameters, and its encoder consists of 24 transformer layers. Only the encoder output is used in this study. For a protein of length *L*, ProtT5 produces a residue-level representation **H**_*P*_∈ ℝ^*L*×1024^after padding and non-residue tokens are removed. Unlike the ESM models, ProtT5 processes long benchmark sequences without any sliding-window procedure. Its self-supervised denoising objective trains the model to reconstruct corrupted amino-acid tokens from the surrounding sequence context.

#### 3) ProstT5

ProstT5 [38] builds on the ProtT5 architecture and pretrained weights and undergoes an additional structure-aware pretraining stage. Protein structures are represented using Foldseek-derived [43] 3Di tokens, and the model learns bidirectional translation between amino-acid sequences and structural-token sequences. ProstT5 retains the 24-layer encoder and 1,024-dimensional residue representation **H**_*P*_ ∈ ℝ^*L×1024*^of ProtT5. During inference, each amino-acid sequence is prepended with the recommended <AA2fold> control token to specify amino-acid-to-structure encoding mode. The control token and padding positions are removed from the final output, yielding **H**_*P*_∈ ℝ^*L×1024*^ after the control token and padding positions are removed. No 3Di tokens or structural coordinates are supplied during benchmark inference, and long sequences are processed without the ESM-specific windowing procedure. ProstT5 is therefore structure-aware through pretraining while remaining sequence-based at inference time.

### C. SELECTED PAIRWISE SIMILARITY METRICS

For every protein pair and every PLM, we compute five pairwise measures between the mean-pooled embeddings 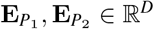 defined in Equations (2) and (3).

#### 1) Cosine Similarity

Cosine similarity uses L2-normalised vectors, so it measures direction only and ignores embedding magnitude:

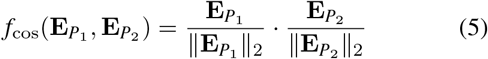

#### 2) Euclidean-Derived Similarity

Euclidean-derived similarity is computed on the raw, unnormalised embeddings:

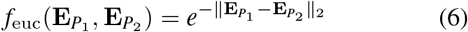

#### 3) Radial Basis Function Similarity

The Radial Basis Function (RBF) kernel similarity applies a Gaussian decay to the squared Euclidean distance:

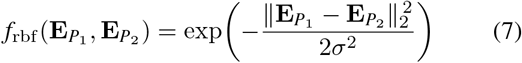

where *σ* is the model-specific RBF bandwidth, set to the median pairwise Euclidean distance computed over all benchmark pairs for that model.

#### 4) Manhattan-Derived Similarity

Manhattan-derived similarity uses the *L*_1_ distance, normalised by the embedding dimension *D*:

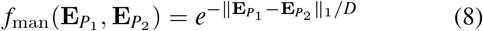

where *D* is the embedding dimension, used to scale the *L*_1_ distance onto a comparable numeric range.

#### 5) Dot Product

The dot product is computed directly on the unnormalised vectors:

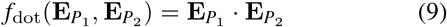

Cosine similarity is the only metric computed on normalised vectors. Equations (6)–(9) are computed on raw, unnormalised embeddings so that each metric remains mathematically distinct from cosine and from one another. This design also allows magnitude-related effects in each model’s embedding space to be observed directly rather than removed by upstream normalisation.

### D. BENCHMARK DATASETS, METHODS TO COMPARE, AND PERFORMANCE EVALUATION

For benchmarking, we use the PISCES dataset [41], containing 20,466 unique protein pairs from 3,500 representative X-ray structures with resolutions ranging from 0.48 to 2.00 Å. All protein pairs share≤ 30% sequence identity, placing the entire dataset within the homology twilight zone [22]. Pairs for which no valid precomputed structural alignment score is available are subsequently excluded, yielding a final benchmark of *N* = 20,445 unique protein pairs [34]. To ensure a fair comparison, the same pair list is used across all PLMs and similarity metrics.

On this dataset, we evaluate embeddings generated by ESM-1b, ESM-2, ProtT5, and ProstT5 using cosine similarity, Euclidean-distance-derived similarity, Manhattan-distance-derived similarity, the radial basis function kernel, and the dot product. To determine whether PLM embeddings identify remote structural relationships beyond those detectable by traditional sequence alignment, we include BLASTP [39], [40] as a sequence-alignment baseline. BLASTP is applied to the same protein pairs, and its log-transformed E-value is used as the primary baseline score. The purpose of including BLASTP is to highlight the performance difference between a traditional approach and alignment-free PLM representations, particularly for remotely related proteins with limited sequence identity.

TM-score, computed using TM-align [42], is used as the ground-truth measure of structural similarity. TM-align requires the three-dimensional coordinates of both proteins, performs a structural superposition, and reports a length-normalised score ranging from 0 to 1. Higher TM-scores indicate better global structural similarity, and values 0.5 or above commonly suggest that two proteins share a similar fold [42]. Because TM-align normalises the structural comparison using each protein length separately, it reports two directional TM-scores for every protein pair. Following prior studies [3], [34], we define TM_min_ as the smaller of the two scores and TM_min_ is used as the primary ground truth in this study.

For each PLM and similarity metric, performance is evaluated using Spearman correlation coefficient, *ρ*, between the embedding-based pairwise similarity scores and TM_min_. To further evaluate performance under different levels of sequence divergence, correlations are calculated separately for the 0% ≤*x*≤ 10%, 10% < *x*≤ 20%, and 20% < *x*≤ 30% sequence-identity intervals. We also evaluate the ability of each method to classify structurally similar protein pairs using TM_min_ as the cutoff, since protein pairs with TMscores (measured by TM-align) greater than or equal to 0.5 are generally considered to share a similar fold [42]. Protein pairs with TM_min_≥ 0.5 are therefore treated as the positive, structurally similar class, whereas those with TM_min_ < 0.5 are treated as the negative, structurally dissimilar class. Receiver operating characteristic (ROC) curves and corresponding area under the ROC curve (AUC) values are calculated across all score thresholds to evaluate each method’s ability to distinguish structurally similar protein pairs from structurally dissimilar pairs. Furthermore, we examine whether sequence-length imbalance affects the performance of PLMs. For each protein pair, the symmetric length ratio is calculated by dividing the length of the shorter sequence by that of the longer sequence. Values close to 1 indicate proteins of comparable length, whereas smaller values indicate greater length imbalance. The benchmark is divided into length-imbalanced pairs with a length ratio≤ 0.5 and more length-balanced pairs with a ratio *>* 0.5, and Spearman correlations with TM_min_ are calculated separately for the two groups to study the impact of sequence-length imbalance on embedding-based similarity performance.

Because the efficiency of our approach depends heavily on generating and storing PLM embeddings, we evaluate the computational cost associated with these two steps. Embedding-generation time is measured on H100 GPU nodes and reported as the average time required by each PLM to generate residue-level embeddings for one protein sequence. We also examine runtime across different sequence-length intervals to determine how embedding-generation time changes with protein length. Storage cost is calculated using the complete set of residue-level embeddings generated for all 3,500 benchmark proteins and represents only the disk space required to store these embeddings. Because storage depends on both the total number of residues and the embedding dimension of each model, this analysis provides a practical comparison of the 1,024-, 1,280-, and 2,560-dimensional representations produced by the evaluated PLMs. We consider residue-level embeddings rather than only the mean-pooled sequence vectors because the complete embedding matrices are typically retained for residue-level analysis, sequence alignment, and other downstream applications. The reported runtime and storage costs do not include similarity calculations or any other downstream processing.

## III. RESULTS AND DISCUSSION

### A. EFFECT OF PLMS AND SIMILARITY METRICS

Table 2 reports Spearman correlation between the embedding-based pairwise similarity scores and TM_min_, the structural ground truth computed using TM-align, across all 20,445 PISCES protein pairs with sequence identity≤ 30%. We evaluate four transformer-based PLMs: ESM-1b, ESM-2, ProtT5, and ProstT5, and use cosine similarity, Euclidean-derived similarity, RBF similarity, Manhattan-derived similarity, and the dot product for each PLM to evaluate the effect of different PLM embeddings and similarity metrics on the performance of protein sequence similarity in the twilight zone. As shown in **Table 2**, ProstT5 with cosine similarity achieves the highest overall Spearman correlation with TM_min_, reaching *ρ* = 0.676. Moreover, ESM-2 (with cosine similarity) provides the next-highest correlation at 0.623, followed by ProtT5 (with cosine similarity) at 0.557 and ESM-1b (with cosine similarity) at 0.483. Moreover, across all five similarity metrics, ProstT5 consistently achieves the highest correlation among the evaluated PLMs, further demonstrating the benefit of its structure-aware pretraining. It is worth mentioning that ProstT5 is the only model exposed to explicit structure-derived 3Di tokens during pretraining; the remaining models are trained on sequence alone. Moreover, the performance improvement over ProtT5 (with cosine similarity), from which ProstT5 inherits both architecture and initial weights, is 0.119, illustrating the effectiveness of the structure-aware pretraining stage rather than any architectural difference. It is also noted that no structural information is supplied at inference time, indicating that structural signal acquired during pretraining is retained in the sequence-derived embedding space.

Furthermore, cosine similarity is the best-performing metric for all four PLMs, indicating that it is the most reliable metric for extracting structure-relevant information from the evaluated sequence-level embeddings. In particular, for ProstT5, the best-performing model, cosine similarity reaches 0.676 and leads the Euclidean-derived and RBF measures at 0.653, Manhattan-derived similarity at 0.648, and the dot product at 0.491. ProtT5 and ESM-1b show a similar pattern, with cosine similarity outperforming other ESM-2, the second-best model, shows a different profile.

**TABLE 2.** Spearman correlations between embedding-based pairwise similarity scores and TM_min_ (measured by TM-align) for 20,445 protein pairs from the PISCES dataset with less than ≤30% sequence identity. Sequence-level embeddings from ESM-1b, ESM-2, ProtT5, and ProstT5 are compared using cosine similarity, Euclidean-derived similarity, the radial basis function (RBF) kernel, Manhattan-derived similarity, and dot product. Higher correlation values indicate stronger agreement with TM_min_. The highest correlation for each PLM is shown in bold.

| PLM | Cosine<br>(Eq. 5) | Euclidean<br>(Eq. 6) | RBF<br>(Eq. 7) | Manhattan<br>(Eq. 8) | Dot Product<br>(Eq. 9) |
| --- | --- | --- | --- | --- | --- |
| ProstT5 | **0.676** | 0.653 | 0.653 | 0.648 | 0.491 |
| ESM-2 | **0.623** | 0.166 | 0.617 | 0.616 | 0.044 |
| ProtT5 | **0.557** | 0.491 | 0.491 | 0.492 | 0.457 |
| ESM-1b | **0.483** | 0.475 | 0.475 | 0.470 | 0.333 |

Cosine similarity attains a strong correlation of 0.623, and RBF and Manhattan-derived similarities remain competitive at 0.617 and 0.616, but the Euclidean-derived measure sharply drops to 0.166 and the dot product to 0.044. Since all five scores are computed from the same ESM-2 embeddings, these differences show that structural-similarity performance is not determined by the PLM alone; it also depends strongly on how the embedding space is compared. Cosine similarity evaluates only the angular relationship between two embeddings. In contrast, the dot product and unnormalized distance measures remain sensitive to embedding magnitude.

**Fig. 2** further shows the joint distribution of cosine similarity of embeddings for each PLM and TM_min_. ESM-1b and ESM-2 concentrate most protein pairs within a narrow cosine-similarity range of approximately 0.85–1.00, despite a broad distribution of TM_min_ values. This compressed cosinesimilarity distribution limits the effective dynamic range available for distinguishing protein pairs. In contrast, ProtT5 distributes pairs across the full interval from 0 to 1, and ProstT5 spans roughly 0.25 to 1.00. This broader dynamic range may facilitate threshold-based discrimination by providing greater separation among protein pairs.

**FIGURE 2.**
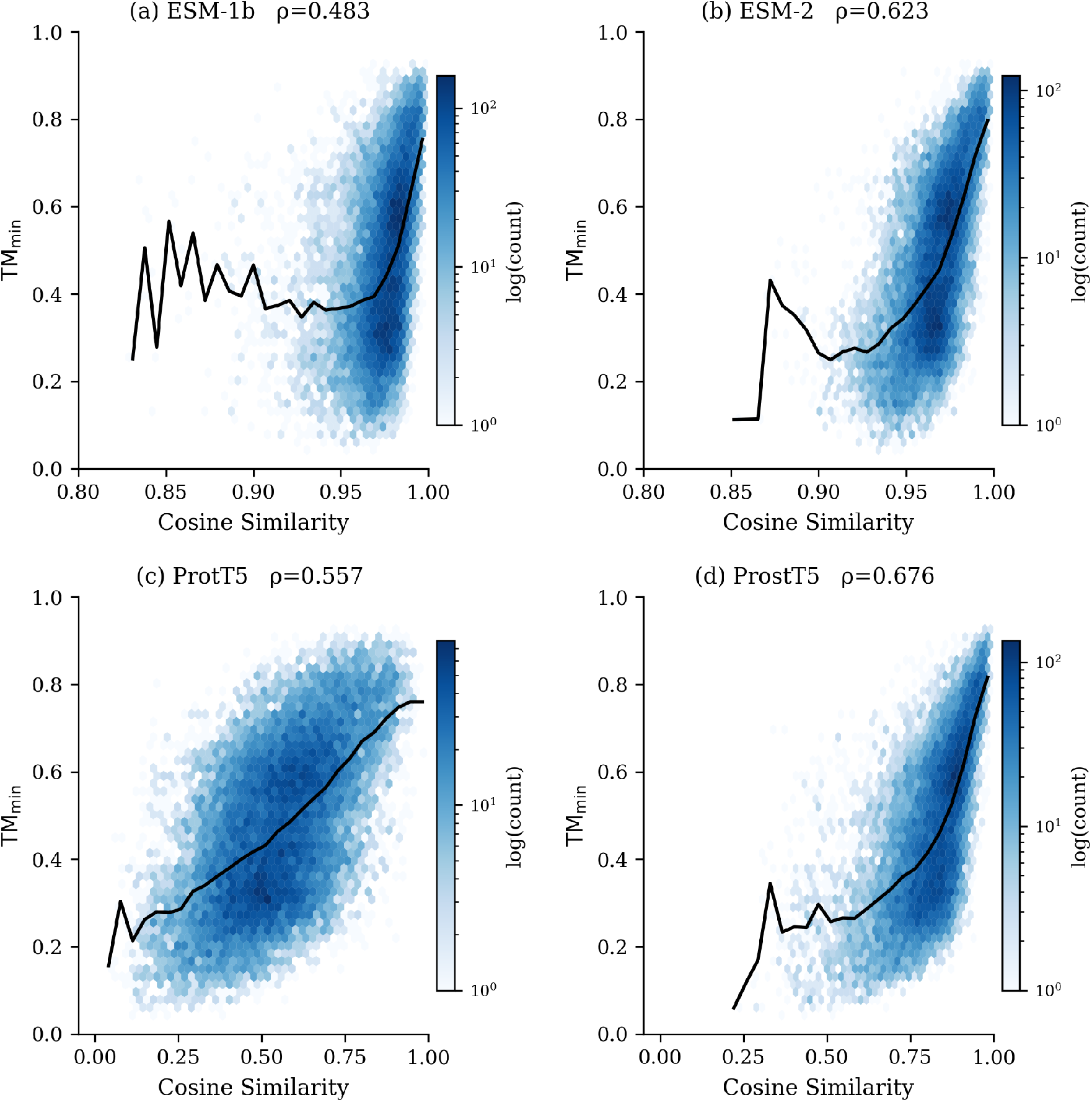
Relationship between cosine similarity and TM_min_ (measured by TM-align) for the four evaluated protein language models across 20,445 PISCES protein pairs with sequence identity≤ 30%. Each panel shows the joint density of TM_min_ and the cosine similarity between mean-pooled sequence embeddings from ESM-1b, ESM-2, ProtT5, and ProstT5. Darker regions indicate a higher density of protein pairs, and the Spearman correlation coefficient (*ρ*) is reported in each panel.

Overall, these results show ProstT5 as the best-performing PLM for protein sequence similarity in the twilight zone and cosine similarity as the most reliable metric across all four PLMs. Cosine similarity is therefore used in all subsequent analyses.

### B. PERFORMANCE COMPARISON AGAINST BLASTP

#### 1) Overall Performance on the PISCES Dataset

**Fig. 3** compares the performance of cosine similarity computed from the mean-pooled embeddings of four transformer-based PLMs with that of BLASTP, a traditional protein sequence alignment tool, by reporting their Spearman correlations with TM_min_ across the same 20,445 protein pairs. BLASTP, acting as a baseline, attains a Spearman correlation of 0.393 with TM_min_, and all four PLMs exceed this baseline. In particular, ESM-1b, the weakest of the four, reaches a Spearman correlation of 0.483 with TM_min_, an improvement of 0.090 over the baseline. ProtT5 reaches 0.557 and ESM-2 reaches 0.623, corresponding to gains of 0.164 and 0.230 over the baseline, respectively. The largest margin is obtained with ProstT5, which reaches a Spearman correlation of 0.676 with TM_min_ and exceeds the baseline by 0.283. It is also worth mentioning that BLASTP fails for 1,410 of the 20,445 protein pairs, whereas each PLM generates an embedding-based similarity score for every pair. Overall, these results demonstrate that mean-pooled PLM embeddings capture structural relationships in the twilight zone more effectively than traditional local sequence alignment.

**FIGURE 3.**
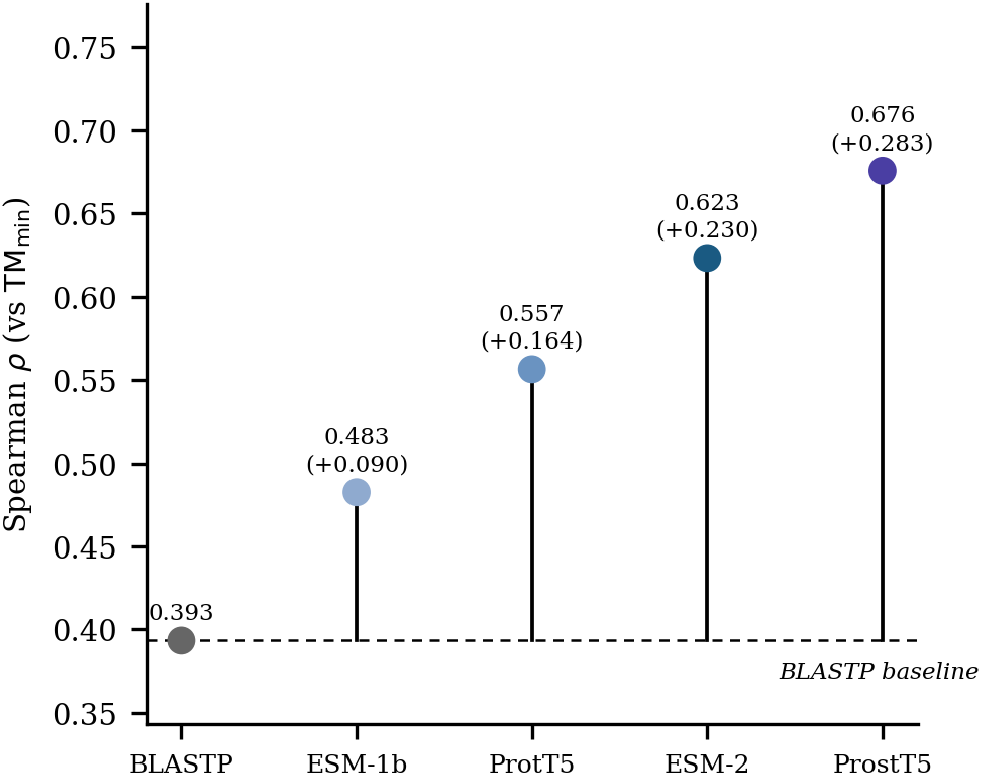
Overall performance comparison of PLM embedding-based similarity and BLASTP (acting as a baseline) using Spearman correlation with TM_min_ across 20,445 PISCES protein pairs with sequence identity≤ 30%. The plotted values show the Spearman correlation between TM_min_ and cosine similarity derived from the mean-pooled embeddings of ESM-1b, ESM-2, ProtT5, and ProstT5, together with the transformed BLASTP score used as the baseline. Values in parentheses indicate the absolute improvement in Spearman correlation over BLASTP.

#### 2) Effect of Sequence Identity

We further evaluate performance of PLM embedding-based similarity and BLASTP by splitting the PISCES dataset into three sequence-identity bins: 0%≤ *x*≤ 10% (*n* = 1,382), 10% < *x*≤ 20% (*n* = 14,671), and 20% < *x*≤ 30% (*n* = 4,392), containing 1,382, 14,671, and 4,392 protein pairs, respectively.

As shown in **Fig. 4**, the Spearman correlations with TM_min_ generally increase with sequence identity for both PLM-based similarity and BLASTP. In the most difficult 0–10% identity interval, ProstT5 achieves the highest correlation at *ρ* = 0.561, followed by ESM-2 at 0.470, ProtT5 at 0.468, and ESM-1b at 0.389. In contrast, BLASTP is nearly uninformative in this interval, reaching a correlation of only 0.019. At 10–20% sequence identity, BLASTP improves to 0.274 but remains below all four PLMs, which reach 0.620 for ProstT5, 0.553 for ESM-2, 0.487 for ProtT5, and 0.433 for ESM-1b. The performance gap over BLASTP remains substantial at 0.346 for the best-performing PLM, ProstT5, and 0.279 for the second best, ESM-2. At 20–30% identity, BLASTP reaches a correlation of 0.726 and becomes considerably more competitive, outperforming both ESM-1b at 0.591 and ProtT5 at 0.689. The two best-performing PLMs, ProstT5 and ESM-2, reach correlations of 0.808 and 0.780, respectively, continuing to lead, but the performance gap drops to 0.082 for ProstT5 and 0.054 for ESM-2. At the same time, the gap between ProstT5 and ESM-2 narrows from 0.091 in the 0–10% interval to 0.067 and 0.028 in the 10–20% and 20–30% intervals, respectively, while ProstT5 remains the best-performing model throughout and shows its largest advantage in the most challenging range. Overall, the result shows that embedding-based similarity provides its greatest advantage over BLASTP at very low sequence identity and also highlights the value of ProstT5’s structure-aware pretraining for highly divergent protein pairs.

**FIGURE 4.**
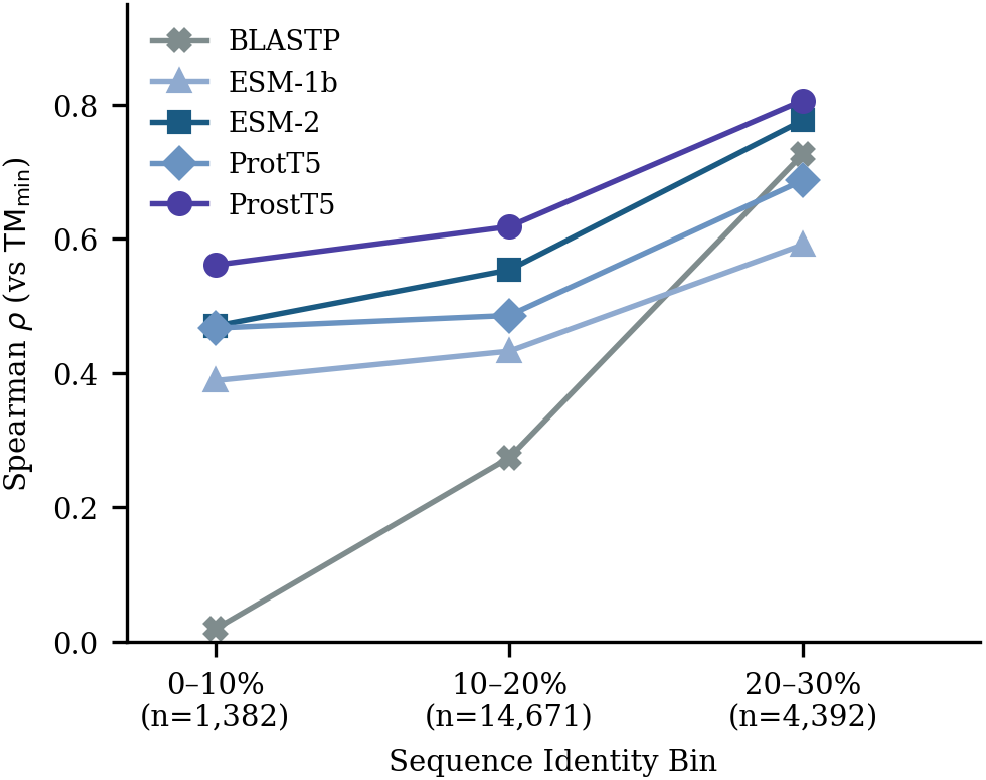
Effect of sequence identity on protein sequence similarity in the twilight zone using PLM embeddings and BLASTP. Protein pairs are divided into three sequence-identity intervals: 0%≤ *x* ≤10% (*n* = 1,382), 10% < *x*≤ 20% (*n* = 14,671), and 20% < *x*≤ 30% (*n* = 4,392). Each point shows the Spearman correlation between TM_min_ and the cosine similarity obtained from mean-pooled ESM-1b, ESM-2, ProtT5, and ProstT5 embeddings. The transformed BLASTP score is included as the baseline. Higher correlation values indicate stronger agreement with structural similarity.

#### 3) Effect of Structural Similarity Threshold

We further assessed whether the evaluated methods can distinguish protein pairs that share a similar fold from those with different folds by using TM_min_ = 0.5 as the structural-similarity threshold. Protein pairs with TM_min_≥ 0.5 were treated as the similar-fold class, yielding 8,998 positive and 11,447 negative pairs, and receiver operating characteristic (ROC) analysis was performed across all possible score thresholds. Cosine similarity is used as the predicted similarity metric for each PLM, whereas the transformed BLASTP log E-value is used for the sequence-alignment baseline. As shown in **Fig. 5**, ProstT5 achieves the best performance in distinguishing structurally similar protein pairs from structurally dissimilar pairs, with an area under the ROC curve (AUC) of 0.831, followed by ESM-2 at 0.806, ProtT5 at 0.766, and ESM-1b at 0.744, whereas BLASTP reaches an AUC of 0.681. Overall, these results show that PLM-based cosine similarity distinguishes structurally similar from structurally dissimilar protein pairs more effectively than BLASTP in the twilight zone, with ProstT5 achieving the best classification performance.

**FIGURE 5.**
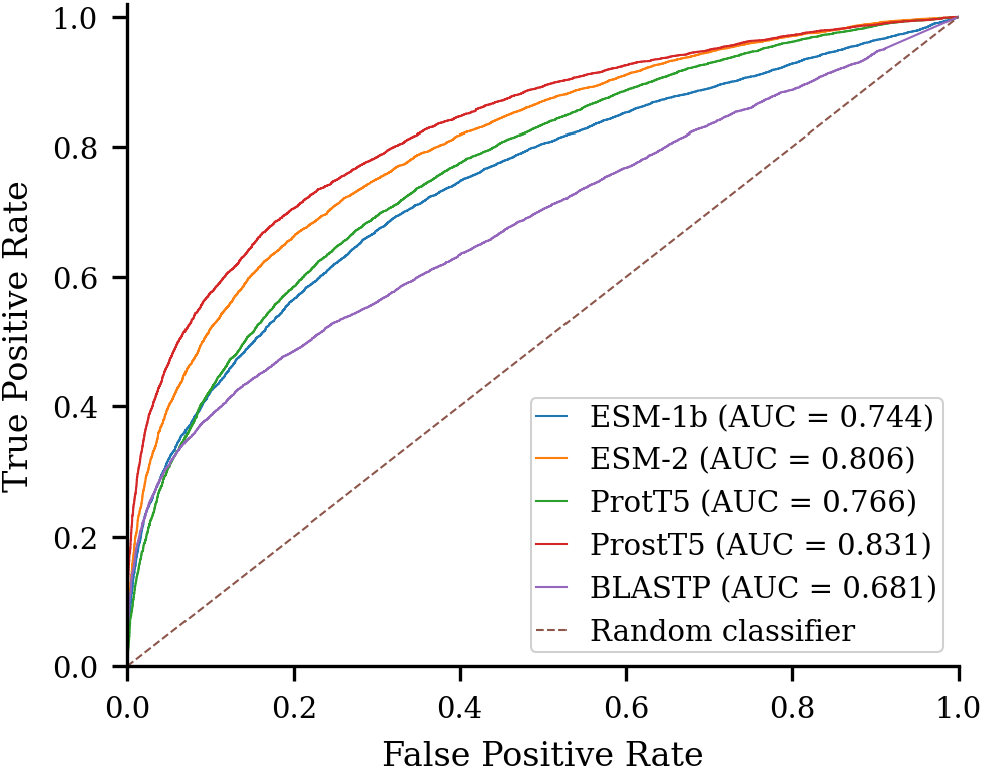
Classification of structurally similar protein pairs using PLM embedding similarities and BLASTP in the protein-sequence twilight zone. Protein pairs with TM_min_ ≥0.5 are treated as the positive, structurally similar class, whereas pairs with TM_min_ < 0.5 are treated as the negative, structurally dissimilar class. ROC curves plot the true-positive rate against the false-positive rate, with AUC values reported in parentheses. Cosine similarities derived from ESM-1b, ESM-2, ProtT5, and ProstT5 embeddings are compared with the transformed BLASTP score. The diagonal dashed line represents random classification with an AUC of 0.5.

These results collectively demonstrate that PLM-based cosine similarity provides a reliable, alignment-free alternative to BLASTP in the twilight zone, with ProstT5 consistently leading across all evaluated sequence-identity ranges and structural-similarity thresholds.

### C. EFFECT OF PAIRWISE SEQUENCE LENGTH RATIO

We next examine how pairwise sequence-length ratio affects the agreement between PLM embedding-based cosine similarity and TM_min_. The symmetric length ratio was calculated by dividing the length of the shorter sequence by that of the longer sequence, and the protein pairs were divided into strongly length-imbalanced pairs with a ratio of 0–0.5 (2,837 pairs) and more length-balanced pairs with a ratio of 0.5–1.0 (17,608 pairs).

As shown in **Fig. 6**, the Spearman correlation with TM_min_ is higher in the more length-balanced group for all four PLMs. ESM-1b shows the largest improvement, increasing by 0.152, from 0.294 to 0.446, followed by ESM-2, which improves by 0.116, from 0.468 to 0.584. ProtT5 and ProstT5 are less affected by sequence-length imbalance, showing smaller improvements of 0.027 and 0.028, respectively. ProstT5 remains the best-performing model in both groups, indicating that its structure-aware representations are comparatively robust to differences in sequence length. Overall, these results suggest that protein sequence similarity obtained from global mean-pooled PLM embeddings is influenced by sequence-length imbalance, with PLM-based protein sequence similarity generally becoming less reliable when one protein is substantially shorter than the other, and ProstT5 and ProtT5 showing greater robustness than ESM-1b and ESM-2.

**FIGURE 6.**
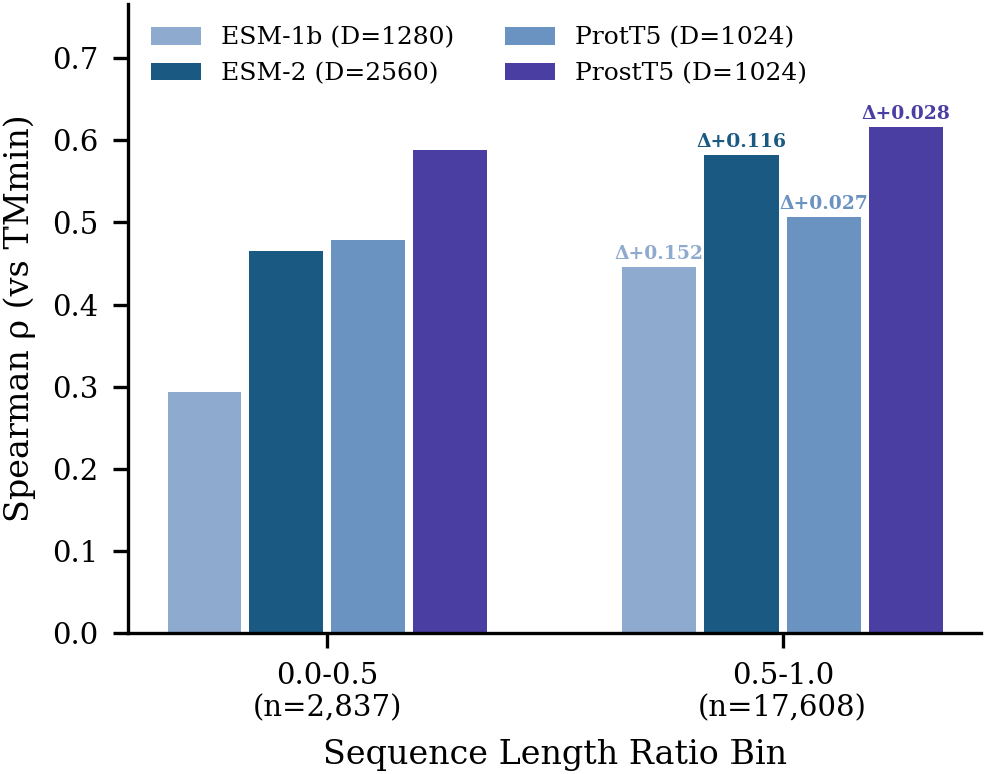
Effect of pairwise sequence-length ratio on protein sequence similarity in the twilight zone. Protein pairs are grouped by symmetric length ratio, defined as the shorter sequence length divided by the longer sequence length, into length-imbalanced pairs (≤ 0.5; *n* = 2,837) and more length-balanced pairs (*>* 0.5; *n* = 17,608). Bars show the Spearman correlation between TM_min_ and cosine similarity computed from the mean-pooled embeddings of each PLM. Values above the second group indicate the absolute increase in correlation relative to the length-imbalanced group, and *D* denotes the embedding dimension.

### D. COMPUTATIONAL EFFICIENCY OF RESIDUE-LEVEL EMBEDDING GENERATION AND STORAGE

We compare the time required to generate residue-level embeddings and the storage required to retain them for the four PLMs. These measurements focus specifically on residue-level embedding generation and storage and do not include mean pooling, similarity calculations, or other downstream processing. It is noted that ESM-2 uses 36 transformer layers and produces a 2,560-dimensional representation per residue, whereas ESM-1b uses 33 layers at 1,280 dimensions, and the two T5 encoders use 24 layers at 1,024 dimensions. **Fig. 7**(a) reports the average residue-level embedding-generation time per protein with respect to protein sequence length on H100 GPU nodes. ESM-1b is the fastest model overall, followed by ProtT5 and then ProstT5; the three are closely comparable, differing by a small margin at every length interval. ESM-2 is clearly the most expensive of the four, and its residue-level embedding-generation time increases considerably more steeply with sequence length. Overall, the best-performing PLM in terms of Spearman correlation with TM_min_, ProstT5, remains close in residue-level embedding-generation time with ESM-1b and ProtT5, whereas the second-best, ESM-2, becomes increasingly expensive as sequence length increases. Moreover, **Fig. 7**(b) compares the total storage required for the complete residue-level embedding collection of all 3,500 benchmark proteins. ProtT5 requires the least storage at 3.08 GB, followed closely by ProstT5 at 3.20 GB. ESM-1b requires 3.85 GB, whereas ESM-2 requires 7.71 GB. Thus, the ESM-2 embeddings require more than twice the storage of the T5-based representations, primarily because of their 2,560-dimensional residue-level output. Overall, ProstT5 offers the most favorable overall trade-off: it achieves the highest correlation with TM_min_ while requiring residue-level embedding-generation time comparable to the fastest models and maintaining one of the smallest residue-level embedding storage requirements. In contrast, ESM-2, the second-most accurate model, is the most expensive on both axes, requiring substantially more time to generate residue-level embeddings for longer sequences and 2.4 times the storage of ProstT5.

**FIGURE 7.**
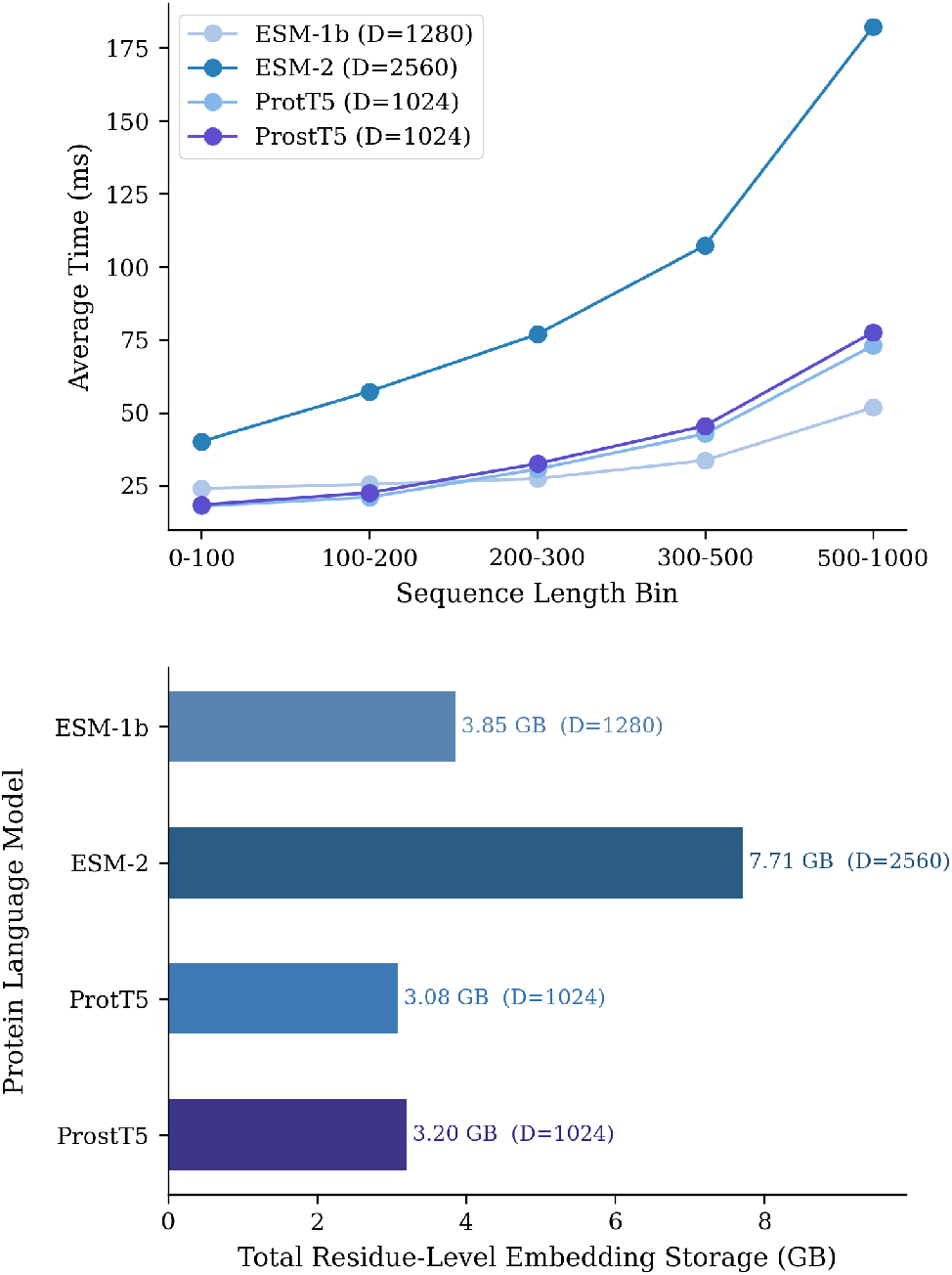
Computational efficiency of residue-level embedding generation and storage for the evaluated PLMs. (a) Average embedding-generation time per protein sequence across five sequence-length bins for ESM-1b, ESM-2, ProtT5, and ProstT5. (b) Total storage required for the residue-level embeddings of all 3,500 PISCES protein sequences. *D* denotes the embedding dimension. Both measurements were obtained using NVIDIA H100 GPU nodes

## IV. CONCLUSION

Accurately identifying protein similarity in the twilight zone remains a major challenge in computational biology. This study evaluates whether sequence-level embeddings from ESM-1b, ESM-2, ProtT5, and ProstT5 can be used to estimate pairwise structural similarity among highly divergent proteins without performing sequence alignment. The evaluation is conducted on the PISCES dataset, which contains protein pairs with sequence identity ≤30% and is therefore suitable for studying protein similarity in the twilight zone, with TM-align-derived TM_min_ used as the structural ground truth. Across the evaluated similarity metrics, cosine similarity provides the most consistent agreement with the structural ground truth, suggesting that the angular relationships between protein embeddings preserve structure-relevant information more effectively than differences in embedding magnitude. Moreover, all four PLMs outperform the BLASTP baseline overall, with the advantage being most evident for protein pairs with very low sequence identity. ProstT5 consistently performs best across the correlation, sequence-identity, and structural-similarity analyses, highlighting the benefit of structure-aware pretraining. The results also show that sequence-length imbalance can reduce the reliability of global embedding similarity, although the T5-based models are comparatively less affected. In addition, ProstT5 provides a favorable balance between similarity performance and the time and storage required to generate and retain residue-level embeddings. Overall, these findings support pretrained PLM embeddings as an effective and practical alignment-free approach for identifying structural relationships among highly divergent proteins using only amino-acid sequences. A limitation of this study is the use of global mean pooling and a single representation layer. Future work will investigate alternative pooling strategies, layer-wise and residue-level representations, broader structural benchmarks, and more recent PLMs, including Ankh.

## ACKNOWLEDGMENT

This work is partially supported by the NSF grant 2435093 (to SB and LW). This work was made possible in part by a grant of high-performance computing resources and technical support from the Alabama Supercomputer Authority.

---

**SHIVARAM DANWADA** received the B.S. degree in computer science, with a specialization in artificial intelligence and machine learning, from Vardhaman College of Engineering, Hyderabad, India, in 2024. He is currently pursuing the M.S. degree in computer science at Auburn University at Montgomery (AUM), Montgomery, AL, USA. His current research interests include applying deep learning to real-world problems, natural language processing, bioinformatics, and AI-based media forensics.

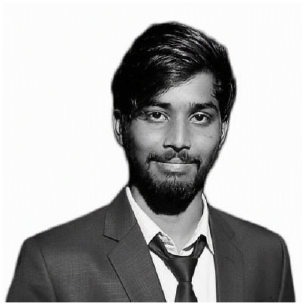

**PRISCILLA UDOMPRASERT** is a recent graduate of Auburn University at Montgomery (AUM) and earned a B.S. degree in computer science. Under the guidance of Dr. Sutanu Bhattacharya, she has been primarily focusing on aiding research with integrating artificial intelligence into bioinformatics. She is currently pursuing another B.S. degree in electrical engineering at Auburn University.

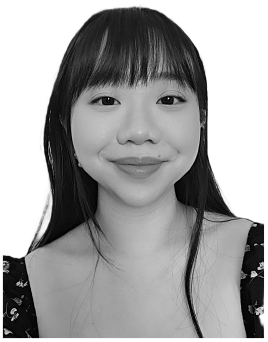

**NILANJANA RAYCHAWDHARY** is currently pursuing the Ph.D. degree in computer science and software engineering with Auburn University, Auburn, AL, USA. She has extensive teaching experience as a Graduate Teaching Assistant and a former Assistant Professor in India. She has actively contributed to diversity in tech through conference talks, awards, and publications. Her research focuses on advancing sentiment analysis in low-resource African languages, such as Igbo, Hausa, and Amharic, using transformer-based models. With multiple peer-reviewed publications, her work addresses critical gaps in NLP for these languages. She was a recipient of the AnitaB.org Advancing Inclusion Scholarship.

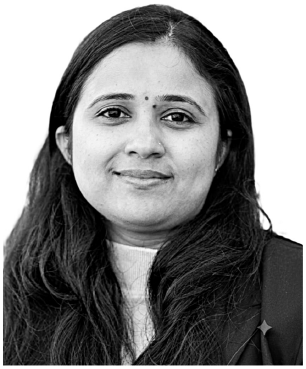

**CHERYL D SEALS** is currently an Associate Professor with the Department of Computer Science and Software Engineering, Auburn University. She also works with outreach initiatives to improve computer science education at all levels. The programs are focused on increasing the computing pipeline by getting students interested in STEM disciplines and future technology careers. Her research interests include human-computer interaction, user interface design, usability evaluation, and educational gaming technologies.

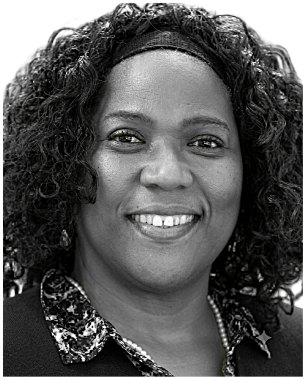

**LEI WU** received his Ph.D. degree in Computer Science from University of Montreal and is currently the Chair and Professor of the Computer Science Department, Auburn University at Montgomery (AUM), Montgomery, AL, USA. His research interests include Artificial Intelligence, Advanced Learning Technology, Robotics Application Development and Autonomous Intelligence.

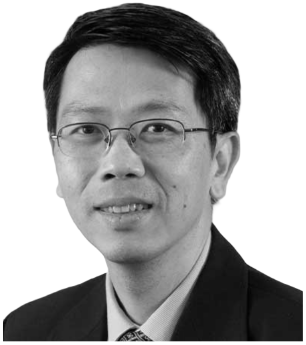

**SUTANU BHATTACHARYA** received his Ph.D. degree in computer science and software engineering from Auburn University, USA. He is currently a tenure-track Assistant Professor with the Department of Computer Science and Computer Information Systems, Auburn University at Montgomery (AUM). Before joining AUM, in 2022, he was an Assistant Professor with the Department of Computer Science, Florida Polytechnic University, Lakeland, FL, USA. His research interests include computational biology, natural language processing, and deep learning.

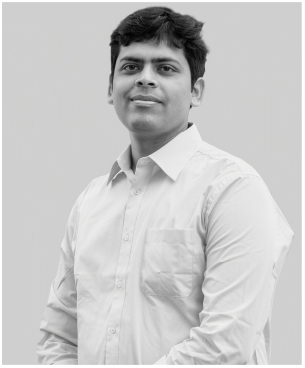

## Notes

This work was partially supported by the National Science Foundation under Grant No. 2435093.

### Competing Interest Statement

The authors have declared no competing interest.

